# ProALIGN: Directly learning alignments for protein structure prediction via exploiting context-specific alignment motifs

**DOI:** 10.1101/2020.12.28.424539

**Authors:** Lupeng Kong, Fusong Ju, Wei-Mou Zheng, Jianwei Zhu, Shiwei Sun, Jinbo Xu, Dongbo Bu

## Abstract

Template-based modeling (TBM), including homology modeling and protein threading, is one of the most reliable techniques for protein structure prediction. It predicts protein structure by building an alignment between the query sequence under prediction and the templates with solved structures. However, it is still very challenging to build the optimal sequence-template alignment, especially when only distantly-related templates are available. Here we report a novel deep learning approach ProALIGN that can predict much more accurate sequence-template alignment. Like protein sequences consisting of sequence motifs, protein alignments are also composed of frequently-occurring alignment motifs with characteristic patterns. Alignment motifs are context-specific as their characteristic patterns are tightly related to sequence contexts of the aligned regions. Inspired by this observation, we represent a protein alignment as a binary matrix (in which 1 denotes an aligned residue pair) and then use a deep convolutional neural network to predict the optimal alignment from the query protein and its template. The trained neural network implicitly but effectively encodes an alignment scoring function, which reduces inaccuracies in the handcrafted scoring functions widely used by the current threading approaches. For a query protein and a template, we apply the neural network to directly infer likelihoods of all possible residue pairs in their entirety, which could effectively consider the correlations among multiple residues. We further construct the alignment with maximum likelihood, and finally build structure model according to the alignment.

Tested on three independent datasets with in total 6,688 protein alignment targets and 80 CASP13 TBM targets, our method achieved much better alignments and 3D structure models than the existing methods including HHpred, CNFpred, CEthreader and DeepThreader. These results clearly demonstrate the effectiveness of exploiting the context-specific alignment motifs by deep learning for protein threading.

## 1 Introduction

Protein tertiary structures are essential to understanding protein functions. Experimental protein structure determination by X-ray crystallography, nuclear magnetic resonance, and cryo-electron microscopy are usually expensive and time-consuming and thus, cannot keep up with the rapid accumulation of protein sequences [25]. Hence, computational approaches to protein structure prediction purely from sequences are highly desirable.

Template-based modeling (TBM), including homology modeling and protein threading, is one of the widely-used methods for protein structure prediction. The rationale underlying TBM is that protein fold types are limited and protein structures are more evolutionarily conserved than protein sequences [15], thus enabling predicting structure of a query protein by referring to structure of homologous proteins. Specifically, for a query protein, TBM first aligns it to all candidate structures (called templates), computes the optimal alignment that maximizes a pre-defined scoring function, and finally constructs 3D structure according to the optimal alignment and corresponding templates [31]. TBM have been very successful in predicting reasonable 3D models for about two thirds of the proteins without solved structures [22].

The performance of TBM relies on extracting informative sequence and structure features from query protein and templates. Most of the existing sequence-template alignment approaches make heavy use of position-specific features, such as sequence profile including position-specific scoring matrix [1] and profile HMM (Hidden Markov Model) [7]. For example, HHpred [26] builds an alignment through the comparison of two profile HMMs. As a residue’s context (i.e., its sequential neighbors) greatly affects its mutation, context-specific features are more informative than position-specific features. CS-BLAST [3] exploits context-specific sequence features, and our previous study [20] exploits context-specific structure features, to obtain more accurate alignments than profile-based alignment approaches. Recently, predicted inter-residue contacts and distances have been proven to be effective in improving sequence-template alignments [34, 33].

The performance of TBM also relies on a scoring function that can accurately measure alignment quality. A successful scoring function should effectively integrate both sequence and structure features. A simple linear combination of protein features is insufficient as most protein features (such as secondary structure and solvent accessibility) are highly correlated [23]. To handle this, the conditional neural field approach uses a probabilistic non-linear function to combine protein features, thus effectively reducing both overcounting and undercounting [18]. DeepThreader [34] and MRFalign [19] use an alignment scoring function that contains singleton terms and pairwise terms. Here, the singleton terms quantify how well a query residue can be aligned with a template residue, while pairwise terms quantify how well two query residues can be aligned with two template residues at a given distance. EigenThreader [4] and CEthreader [33] make use of predicted inter-residue contact map, while DeepThreader makes use of predicted distance map.

In this work, we present a novel approach called ProALIGN for protein alignment and threading. Unlike many existing approaches that build alignments using a handcrafted scoring function, our approach directly learns and infers protein alignments. Our approach is founded on the observation of alignment motifs: Protein sequences consist of sequence motifs with conserved amino acid composition, and protein structures consist of structure motifs with conserved spatial shapes. Similarly, protein alignments also consist of alignment motifs formed by aligned structure motifs. Alignment motifs, which appear frequently in protein alignments, usually exhibit characteristic patterns. As shown in Figure 1, when representing a protein alignment as a matrix with dots denoting aligned residue pairs, aligned helices show as diagonal lines while alignment gaps show as two diagonal lines with a shift between them. In addition, alignment motifs are context-specific as their characteristic patterns are tightly related to sequence contexts of the aligned regions. These observations enable us to recognize alignment motifs based on sequence contexts.

**Fig. 1.**
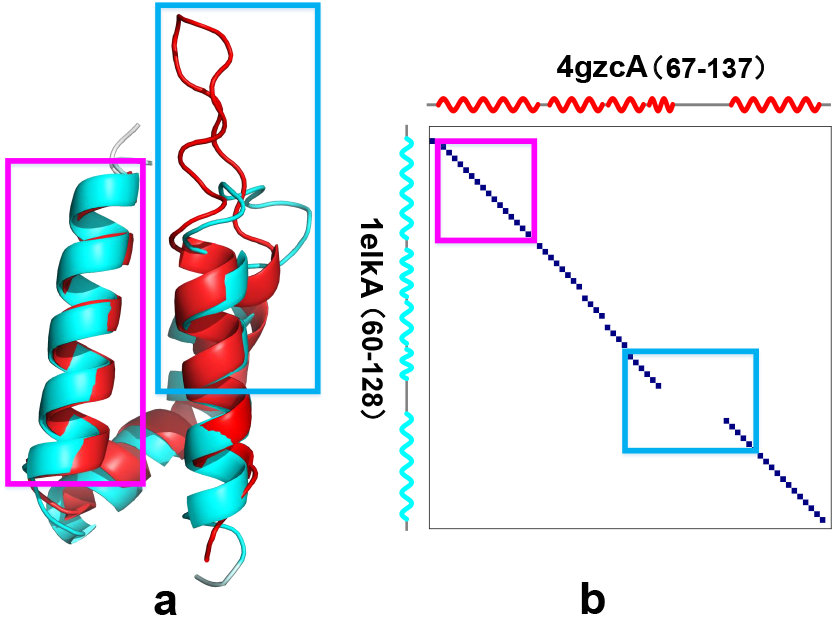
Protein alignment and alignment motifs. (*a*) The structure alignment of protein 4gzcA (in red) and 1elkA (in cyan). Only parts of these proteins are shown here due to the limited space. The rectangle in magenta shows two aligned helices (Y69-V84 of 4gzcA vs. N63-V78 of 1elkA) while the rectangle in blue shows an alignment gap (F99-S114 of 4gzcA vs. T102-S123 of 1elkA). (*b*) The matrix representation of the alignment, where a dot represents an aligned residue pair, highlights alignment motifs with characteristic patterns. For example, the aligned helices shows as a diagonal line (the rectangle in magenta) whereas the alignment gap shows as two diagonal lines with a shift between them (the rectangle in blue)

Deep convolutional network has shown great success in pattern recognition, especially for image processing such as classification and object detection [14,5]. By treating alignment matrices as images, we utilize a deep convolutional neural network to directly learn protein alignments by integrating both sequential information and predicted inter-residue distances. The trained neural network implicitly but effectively encodes an alignment scoring function, which reduces inaccuracies in the handcrafted scoring function widely used by the current threading approaches. For a query protein and a template, we apply the neural network to infer likelihoods of all possible residue pairs in their entirety, which could effectively consider the correlations among multiple residues. We further construct the optimal alignment with maximum likelihood, and finally build structure model according to the alignment.

Using three independent benchmark datasets including 6688 protein alignment targets and 80 CASP13 TBM targets, we show that ProALIGN may produce much more accurate alignments and 3D models than the state-of-the-art approaches including HHpred, CNFpred, CEthreader and DeepThreader.

## 2 Methods

### 2.1 Overall workflow of ProALIGN

The workflow of our ProALIGN approach for TBM is shown in Figure 2. It consists of the following 4 main steps: *(i) Feature calculation:* The features to be used include sequence profile, secondary structure, solvent accessibility, and inter-residue distances. *(ii) Alignment likelihood inference:* The input features are feed into a pre-trained deep convolutional neural network, which predicts alignment likelihood for each residue pair (one query residue and one template residue). In our approach, alignment likelihood are represented as a matrix form. One entry in the matrix contains the match likelihood value of a residue pair. *(iii) Alignment generation:* Based on the alignment likelihood, we construct the optimal alignment with maximum likelihood. *(iv) Model building:* Build 3D structure model by running MODELLER [8] on the generated alignment.

**Fig. 2.**
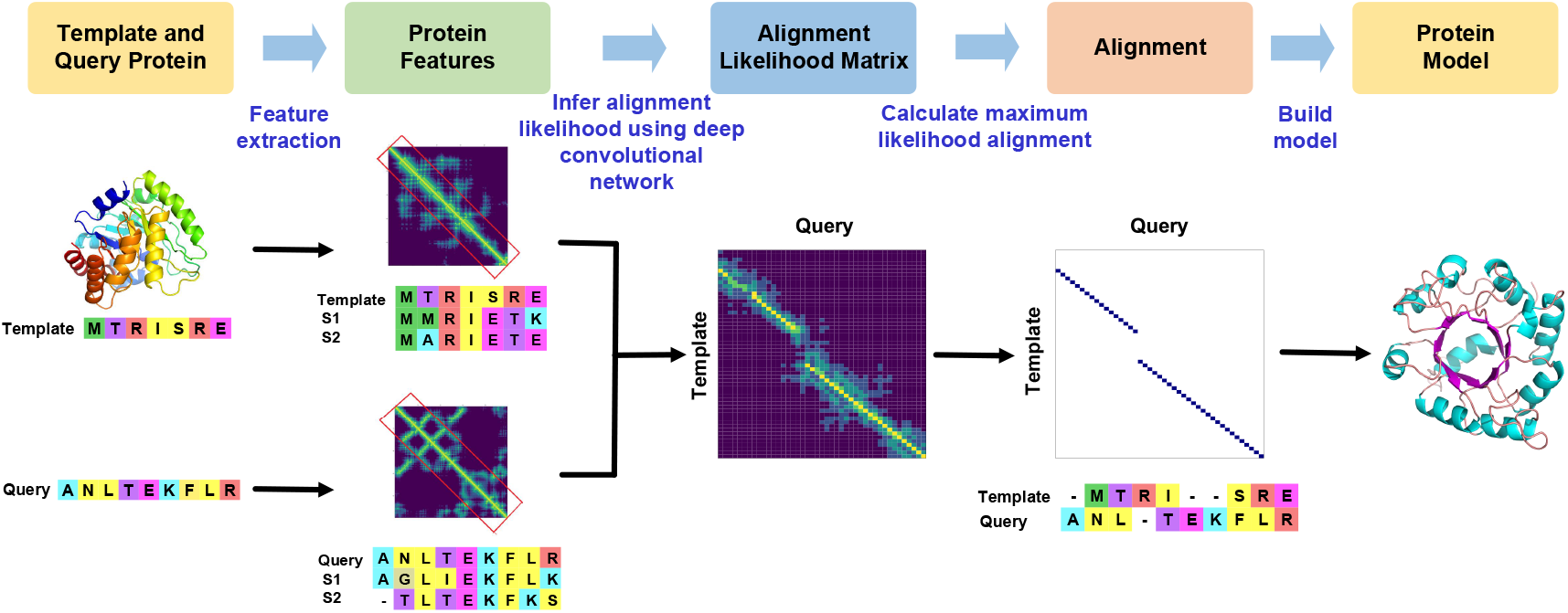
The overall workflow of ProALIGN for TBM. It has 4 main steps: *(i) Feature calculation; (ii) Alignment likelihood inference; (iii) Alignment generation;* and *(iv) 3D model building.*

Meanwhile, the first two steps are different from the existing approaches and thus, will be described in detail very soon.

### 2.2 Representation of alignment and formulation of alignment problem

For a query protein with *n* residues *S* = *S*_1_*S*_2_…*S_n_* and a template with *m* residues *T* = *T*_1_*T*_2_…*T_m_,* we represent an alignment of them as a binary matrix *A*:

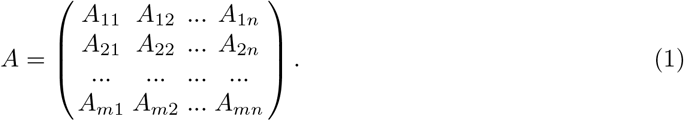

Here, *A_ij_* = 1 if and only if template residue *T_i_* is aligned with query residue *S_j_* (Fig. 3). An alignment matrix can be intuitively treated as an image and thus, amenable to deep convolutional neural networks, which are good at learning patterns in the alignment matrix.

**Fig. 3.**
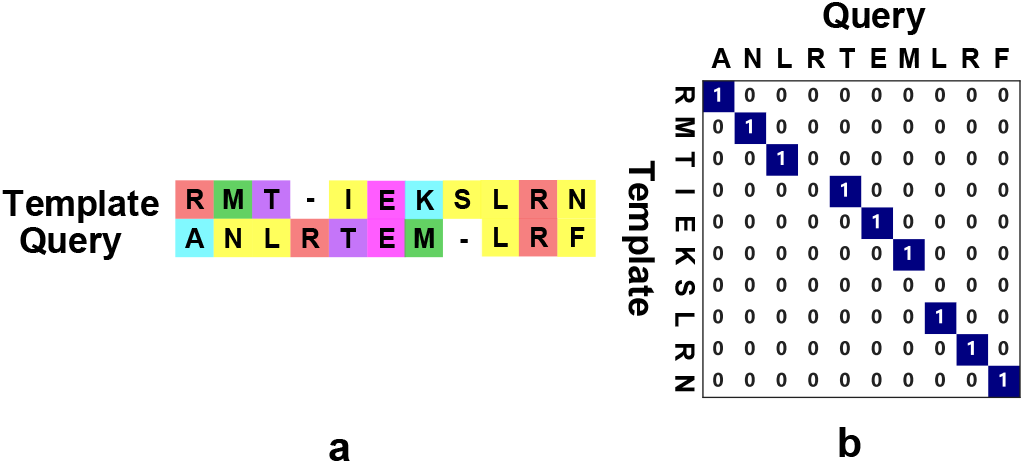
Two equivalent representations of a protein alignment. (*a*) Representation of an alignment as a sequence pair with symbol ‘-’ denoting gaps. (*b*) Matrix representation. Intuitively, an alignment matrix can be treated as an image and thus, suitable for learning and inference by deep convolutional neural networks.

Given a protein pair *T* and *S*, we use a deep convolutional neural network with parameter *θ* to learn and infer their alignment. The probability of an alignment *A* can be represented as:

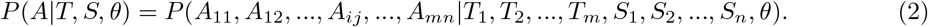

Specifically, for a pair of residues *T*_i_ and *S_j_*, the neural network predicts their likelihood of being aligned (i.e., *A_ij_* = 1). Our neural network simultaneously estimates the likelihood of all the residue pairs being aligned so that it can take into consideration the correlations among residue pairs.

### 2.3 Learning protein alignments using deep convolutional neural network

We design a deep convolutional neural network to learn the relationship between characteristic pattern of alignment motifs and residue composition of the corresponding sequence contexts. Here, the residue composition of sequence contexts are represented using a collection of protein features, which construct the input of our neural network.

#### Input features

Our neural network takes sequence features and structure features of proteins as input. Specifically, for both template and query proteins, we first construct multiple sequence alignment through running HHblits [24] against a sequence database, and then build position-specific scoring matrix (PSSM), which could effectively depict the preference of amino acid types at each position.

The structure features used in this study include secondary structure, solvent accessibility, and inter-residue distances. For a template, secondary structure and solvent accessibility are calculated by running DSSP [9] on the template structure. The inter-residue distances are calculated according to the coordinates of *C_β_* atom of residues. For a query protein, we predict its secondary structure and solvent accessibility using RaptorX-Property [29] and predict inter-residue distance using ProFOLD [11]. For each residue, we use the distance with neighbor residues within a window. Here, we set the window size as *C* =33 (The other settings of window size, e.g., 1-41, were also examined and the prediction results are stable when *C* ≥ 29).

Thus, for each residue in query or template protein, we obtain a total of 356 features, including 20 PSSM features, 3 secondary structure features, 3 solvent accessibility features, and 330 interresidue distance features (33 neighboring residues times 10 distance bins). For each residue pair containing a query residue and a template residue, we concatenate their features, thus obtaining a total of *m* × *n* × 712 features. The features of all residue pairs construct template-query joint features (Fig. 4). Our neural network takes these joint features as input.

**Fig. 4.**
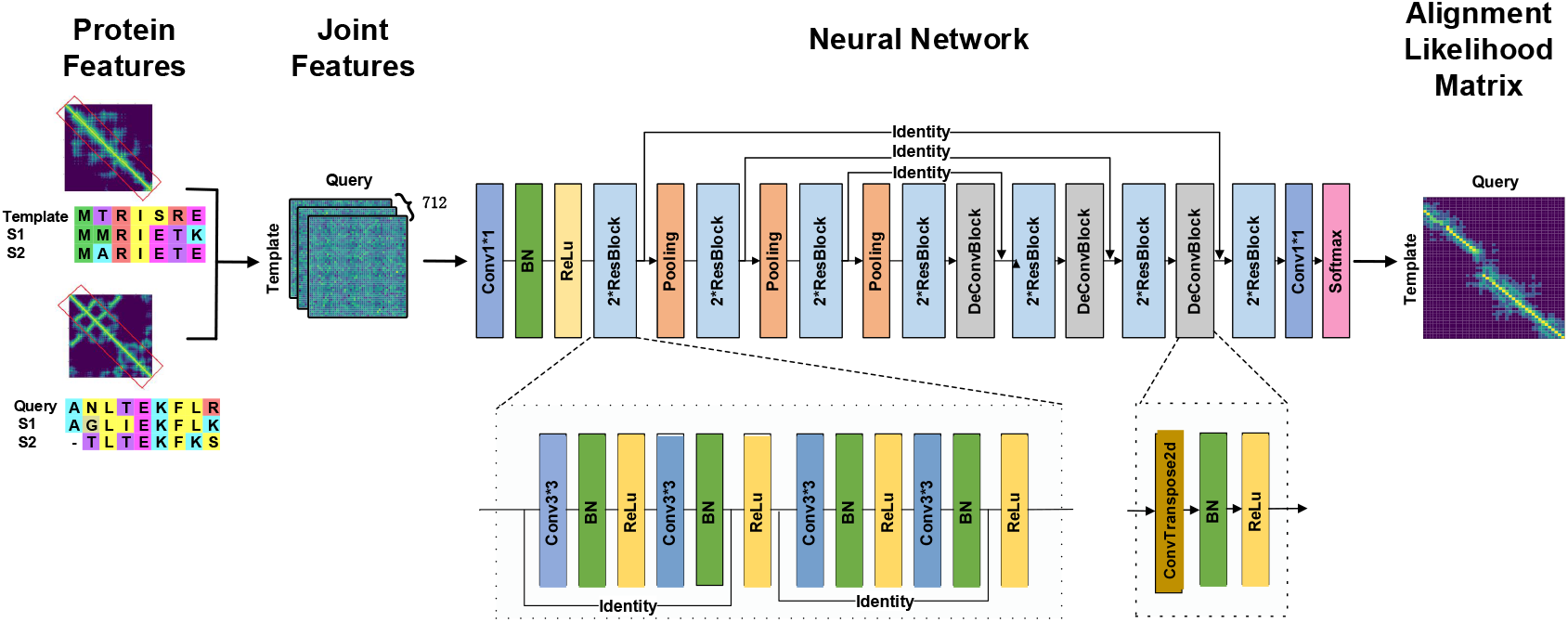
Architecture of the deep neural network used by ProALIGN. The neural network takes sequence features and structure features of proteins as input, and outputs an alignment likelihood matrix in its entirety. The (*i,j*) entry of the matrix represents the likelihood whether template residue *i* assigns with query residue *j.* The network has a multi-layer convolution stack structure. Using this structure, we expect that the shallower layers can capture the characteristic pattern of alignment motifs, and the deeper layers can learn the correlation among alignment motifs

#### Neural network architecture

To learn the relationship between alignment and the joint protein features, we design a deep neural network with full convolutional architecture [16]. Full convolutional architecture does not assume input length to be fixed, and thus can handle proteins with variable lengths. Using multi-layer convolutional stack structure, full convolutional network is expected to capture both coarse and fine information. The shallower convolutional layers have smaller receptive field and learn detailed features of local areas, while the deeper layers have larger receptive field and thus can learn more global and abstract information (Figure 4). Here, we expect that the shallower layers can capture the characteristic pattern of alignment motifs, and the deeper layers can learn the correlation among alignment motifs.

We also add several shortcut connections (identity operations)[16] into the neural network. Using these shortcut connections, we can effectively combine the information captured by shallower and deeper layers. To reduce the risk of gradient vanishing, we use the residual network structure[10]. The whole neural network adopts ReLU[21] as activation function.

#### Loss function

Each sample of the training set used in this study contains two proteins with their structural alignment (called reference alignment) as label. The objective of training process is to find optimal network parameter to maximize the probability that the network generates the given reference alignment. For this end, we use the cross-entropy loss function as follows.

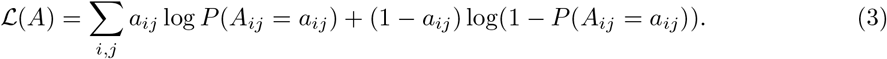

Here, *a_ij_* ∈ {0,1} denotes label for residue pair (*i,j*), i.e., whether residue *i* of template protein aligns with residue j of the query protein in their reference alignment, and *A_ij_* denotes network’s output for these two residues.

#### Training the neural network

We train the neural network using Adam algorithm [12]. To prevent overfitting, we adopt weight decay operation with coefficient of 0.001 during gradient updating.

The neural network is trained using mini-batch[27] technique. As proteins in a batch always have different size, zeroes should be padded to make their sizes equal. However, when a protein has extremely long size, this zero-padding operation might lead to a large memory requirement that exceeds GPU’s capacity. To overcome this difficulty, we apply large batch size for short proteins and small batch size for long proteins.

### 2.4 Inferring protein alignment using deep convolution neural network

We apply the trained neural network on a query protein *S* and a template *T*, and obtain an alignment likelihood matrix. The (*i,j*) entry of the matrix represents the likelihood whether template residue *i* assigns with query residue.

Note that a valid alignment A poses two restrictions on the aligned residue pairs: (*i*) A residue aligns with at most one residue. (*ii*) Two aligned residue pairs cannot crossover, i.e., suppose query residue *i* aligns template residue *j*, and query residue *k* (*k* > *i*) aligns template residue *l*, then *l* should be large than *j*. These restrictions can be intuitively demonstrated from the view point of alignment matrix: there is at most a ‘1’ in one column and one row, and the aligned residue pairs show a shape of diagonal line rather than a back-diagonal line. Here, we denote the set of all valid alignments as *U*. The optimal alignment *A_opt_* is expected to obtained by maximizing the matrix likelihood, at the same time satisfying the definition of valid alignment, i.e.,

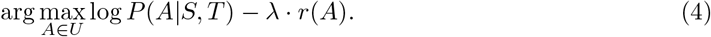

Here, *r*(*A*) represents a regular term to make the alignment compacted. In this study, we set *r*(*A*) as the length of actual alignment area, i.e., the area from the first aligned residue pair to the last aligned residue pair in built alignment. We expected this regular term to penalize too many mismatch residues in alignment. The optimal setting of weight λ is decided using validation dataset Valid1K (described in later). In this study, we apply dynamic programming to solve the optimal alignment that satisfies valid alignment restrictions.

## 3 Results and discussions

### 3.1 Training and test data

#### Training dataset

We used PDB25 dataset (constructed from PISCES[28] in 2015, containing 7,952 proteins in which any two proteins share ≤25% sequence identity) to construct training and test dataset. To guarantee no overlap between training set and testing set, we first divided the PDB25 proteins into two parts: one part containing 6,245 proteins for training and validation, and the other part containing 1,707 proteins for testing. From the first part (containing 6,245 proteins), We calculate the structure alignment of any two proteins as reference alignment using structure alignment tool DeepAlign [30]. After filtering out low-quality alignments (TMscore < 0.40), we obtained a total of 75,874 alignments. From these alignments, we randomly selected 67,106 alignments as training set and used the remainder 8,768 as validation set. In addition, we also randomly select 1000 alignment pairs (Valid1K) to decide the hyper-parameter λ used in Eq. 4.

#### Test dataset

From the second part of PDB25 proteins(containing 1,707 proteins), we generated a total of 5,688 alignments for testing (referred to as *Testing5.6K*). In addition, we also evaluated our approach on the 1,000 protein alignments (referred to as *Testing1K*) used by DeepThreader. After removing the low quality alignments (TMscore < 0.40), we acquired a total of 769 alignments with high quality. According to structure similarity, we split these alignments into three groups, i.e., easy group with TMscore in (0.80, 1], medium group with TMscore in (0.60. 0.80], and hard group with TMscore in (0.40, 0.60].

To evaluate threading performance, we used the 80 domains released by CASP13 organizer, including 22 TBM Hard domains, 45 TBM Easy domains, and 13 FM/TBM domains. We used PDB40 containing 32,363 proteins as template. It is worthy pointed out that the PDB40 was constructed on April 4, 2018 before CASP13, thus avoiding potential misusing the native structure of CASP13 domains as templates.

### 3.2 Evaluation method

To evaluate alignments, we calculated both reference-dependent and reference-independent alignment accuracy. Here, the reference-dependent accuracy is defined as the percentage of correctly aligned positions judged by the reference alignments, and the reference-independent accuracy of an alignment is defined as quality of the 3D model generated from the alignment. Here, we applied MODELLER to generate 3D model from an alignment.

We compared ProALIGN with the state-of-the-art alignment approaches, including HHpred, CNFpred, CEthreader, and DeepThreader. For the sake of fair comparison, we executed these programs with the same MSA (constructed using HHblits with 3 iterations and E-value set to 0.001 against Uniprot20_2016[2]). We run HHpred with mode ‘-mact 0.1’ and run CEthreader with ‘EigenProfileAlign’ mode.

### 3.3 Reference-dependent alignment accuracy

Table 1 shows the reference-dependent alignment accuracy on Test5.6K dataset. To reduce the potential bias in generating reference alignments, we evaluated our approach using reference alignments generated using three structure alignment tools, including TMalign[32], DeepAlign and MM-Ligner[6].

**Table 1.**
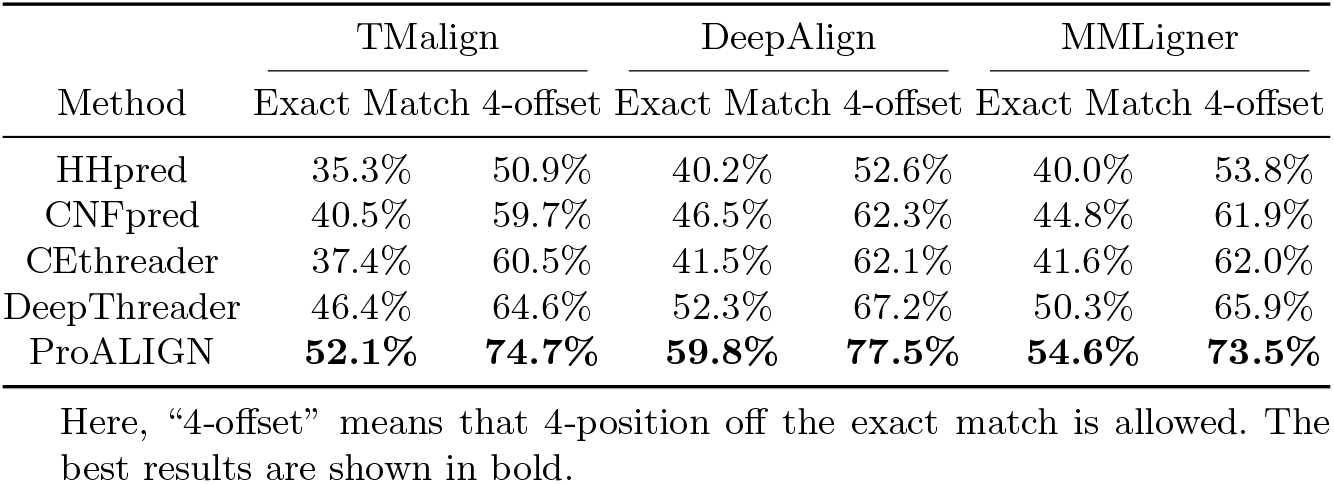
Reference-dependent alignment accuracy on Test5.6K dataset

When using the TMalign reference alignment, ProALIGN shows an alignment accuracy (i.e., exact match) of 52.1%, which outperforms DeepThreader, CEthreader, CNFpred and HHpred by 5.7%, 14.7%, 11.6% and 16.8%, respectively. If 4-position off the exact match is allowed in calculating alignment accuracy, ProALIGN shows an alignment accuracy of 74.7%, higher than DeepThreader, CEthreader, CNFpred and HHpred by 10.1%, 14.2%, 15% and 23.8%, respectively. Similar observations can be obtained when using DeepAlign and MMLigner reference alignment.

As shown in Table 2, the alignment accuracy of ProALIGN is much higher on Test1K dataset (69.0% for exact match, and 85.3% for 4-offset when using TMalign reference alignment). Again, ProALIGN significantly outperforms the existing approaches. These results clearly suggest that ProALIGN can generate more accurate alignments for proteins.

**Table 2.**
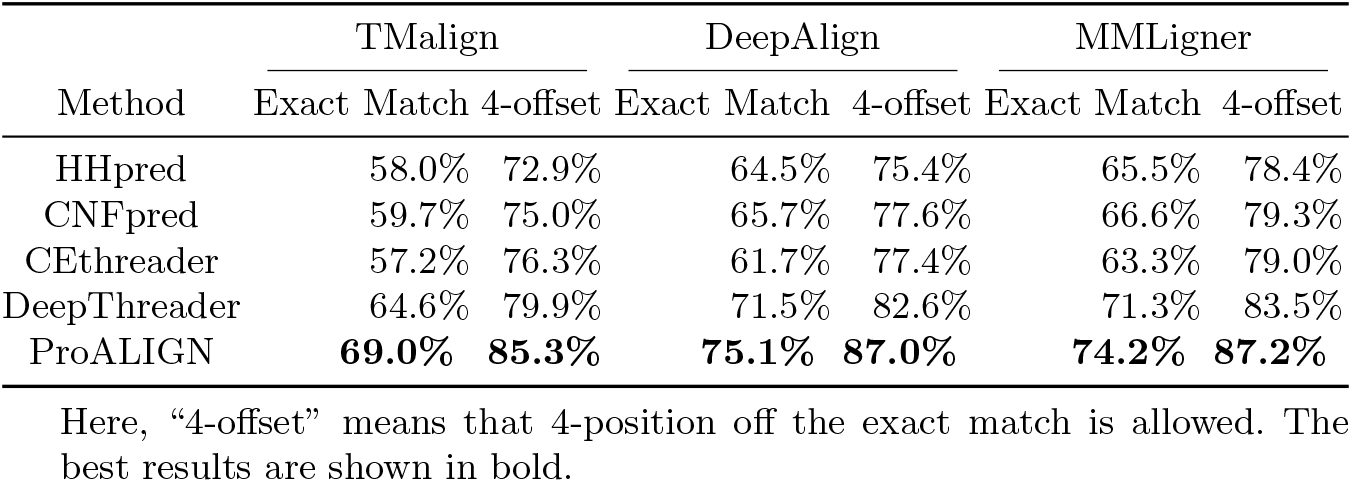
Reference-dependent alignment accuracy on Test1K dataset

### 3.4 Reference-independent alignment accuracy

Further, we assessed ProALIGN and other alignment approaches in terms of reference-independent alignment accuracy. As shown in Table 3, the model built by ProALIGN have an average TMscore of 0.551, which is much better than DeepThreader (0.511), CEthreader (0.470), CNFpred (0.467) and HHpred (0.406). Besides examining the average TMscore of all generated models, we further investigated each model individually using head-to-head analysis. Figure 5 suggests that ProALIGN could generate higher quality models in most cases.

**Fig. 5.**
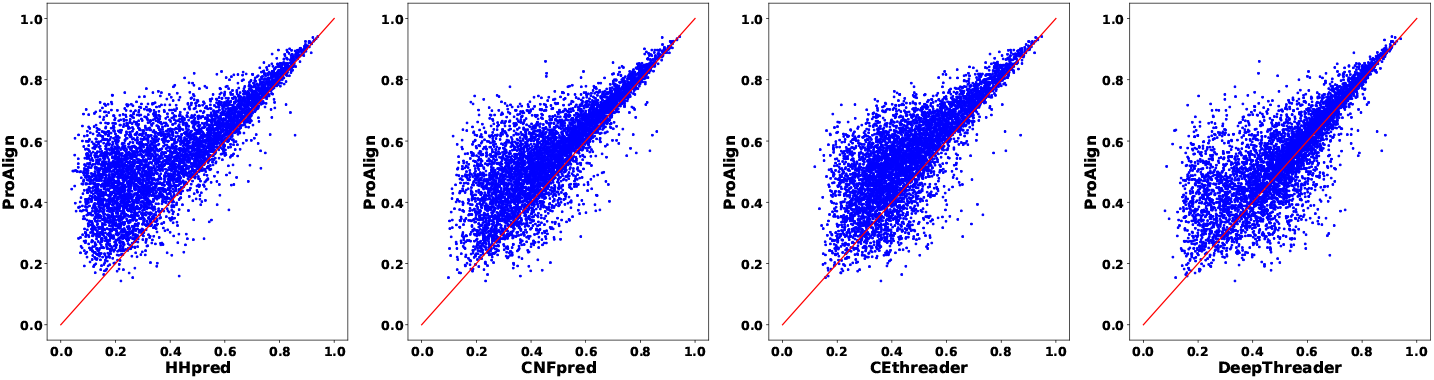
Head-to-head comparison of the quality of models generated using ProALIGN, HHpred, CNpred, CEthreader, and DeepThreader on Test5.6K dataset. Here, the model quality is measured using TMscore

**Table 3.** Reference-independent alignment accuracy measured by TMscore and GDT on Test5.6K

| Group | #Alignments | HHpred |  | CNFpred |  | CEthreader |  | DeepThreader |  | ProALIGN |  |
| --- | --- | --- | --- | --- | --- | --- | --- | --- | --- | --- | --- |
|  |  | TMscore | GDT | TMscore | GDT | TMscore | GDT | TMscore | GDT | TMscore | GDT |
| Test5.6K Easy | 216 | 0.798 | 0.673 | 0.810 | 0.690 | 0.803 | 0.679 | 0.806 | 0.682 | **0.821** | **0.700** |
| Test5.6K Medium | 1971 | 0.533 | 0.440 | 0.580 | 0.486 | 0.578 | 0.481 | 0.614 | 0.513 | **0.653** | **0.550** |
| Test5.6K Hard | 3501 | 0.310 | 0.244 | 0.382 | 0.301 | 0.389 | 0.306 | 0.435 | 0.344 | **0.476** | **0.380** |
| Test5.6K | 5688 | 0.406 | 0.328 | 0.467 | 0.380 | 0.470 | 0.381 | 0.511 | 0.416 | **0.551** | **0.451** |

In particular, for the Test5.6K Easy group, all of these alignment approaches could generate high-quality models. For the Test5.6K Medium group, the superiority of ProALIGN is obvious: the average TMscore of the generated model is 0.653, higher than HHpred (0.533), CNFpred (0.580), CEthreader (0.578) and DeepThreader (0.614). For the Test5.6K Hard group, ProALIGN shows an average TMscore of 0.476, outperforms HHpred, CNFpred, CEthreader and DeepThreader by 0.166, 0.094, 0.087 and 0.041, respectively.

Evaluation on Test1K dataset confirmed the superiority of ProALIGN (Table 4 and Figure 6). These results together suggested that ProALIGN could generate 3D model with higher quality than the existing approaches, especially for the medium and hard protein alignments.

**Fig. 6.**
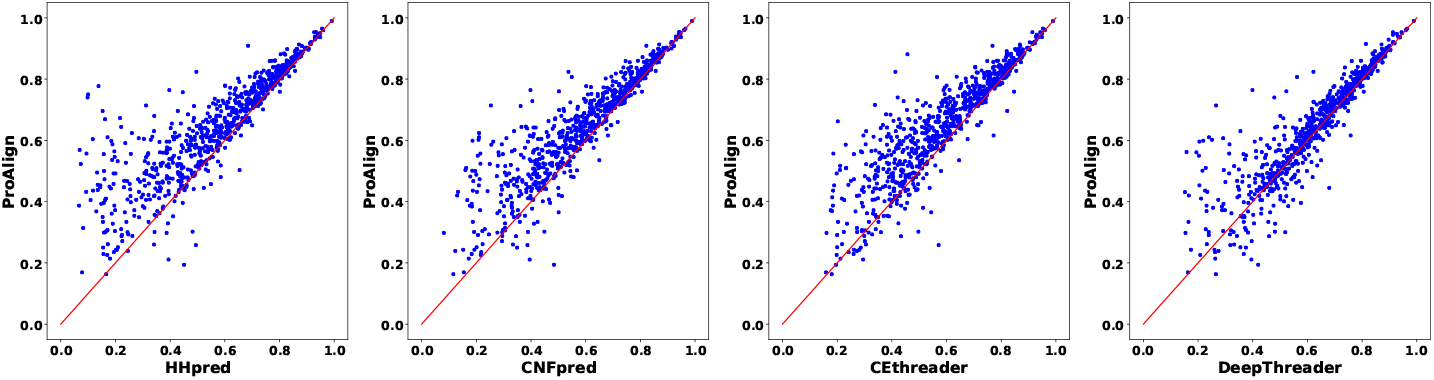
Head-to-head comparison of the quality of models generated using ProALIGN, HHpred, CNpred, CEthreader, and DeepThreader on Test1K dataset. Here, the model quality is measured using TMscore

**Fig. 7.**
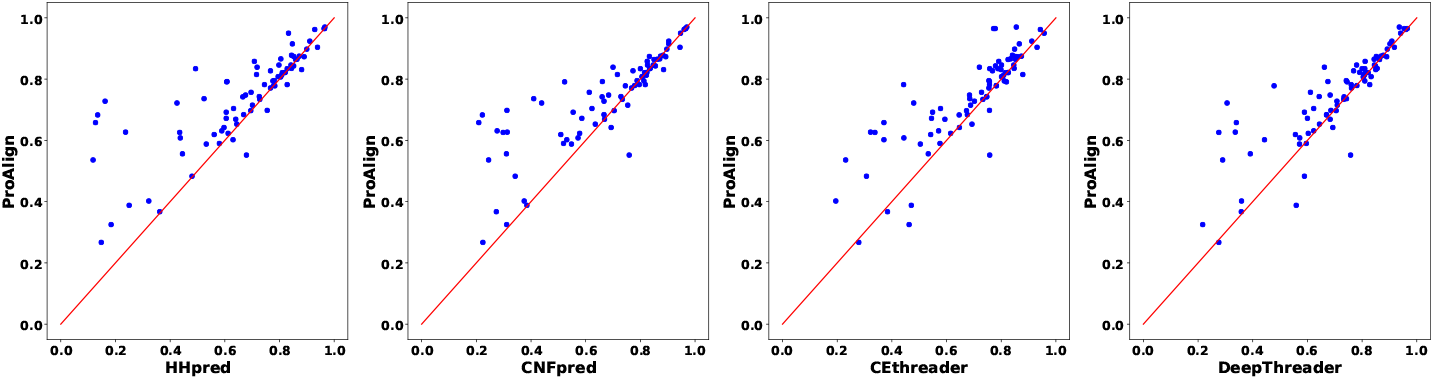
Head-to-head comparison of quality (measured using TMscore) of the top1 model predicted by HHpred, CNFpred, CEthreader, and DeepThreader on 80 CASP13 TBM targets

**Table 4.** Reference-independent alignment accuracy measured by TMscore and GDT on on Test1K

| Group | #Alignments | HHpred |  | CNFpred |  | CEthreader |  | DeepThreader |  | ProALIGN |  |
| --- | --- | --- | --- | --- | --- | --- | --- | --- | --- | --- | --- |
|  |  | TMsore | GDT | TMsore | GDT | TMsore | GDT | TMsore | GDT | TMsore | GDT |
| Test1K Easy | 121 | 0.822 | 0.752 | 0.831 | 0.763 | 0.826 | 0.754 | 0.838 | 0.769 | **0.846** | **0.776** |
| Test1K Medium | 334 | 0.647 | 0.544 | 0.660 | 0.558 | 0.655 | 0.550 | 0.694 | 0.586 | **0.715** | **0.605** |
| Test1K Hard | 314 | 0.395 | 0.314 | 0.418 | 0.333 | 0.418 | 0.330 | 0.472 | 0.375 | **0.497** | **0.398** |
| Test1K | 769 | 0.571 | 0.483 | 0.588 | 0.498 | 0.585 | 0.492 | 0.626 | 0.529 | **0.646** | **0.548** |

### 3.5 Threading performance on CASP13 TBM targets

When using threading approach for protein structure prediction, we need to align the query protein with all templates, and then select the most reliable alignment and template to build 3D model. Thus, a strategy is required to rank alignments.

As performed by CEthreader, we ranked alignments according to the ratio of query contacts that are aligned onto template contacts, i.e., 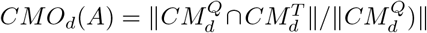. Here, 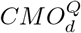 and 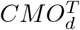 represent the number of residue contacts with distance less than threshold *d* for query protein *Q* and template *T*, respectively. The average of the ratios at four different distance thresholds 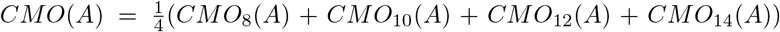 was finally used to rank alignments. For each target, we evaluated two predicted models: (*i*) top1 model: the model built based on the alignment ranked 1st, and (*ii*) top5 model: the best model among the models built based on top 5 alignments.

Table 5 shows quality (measured using TMscore and GDT) of the top1 and top5 model predicted for the CASP13 TBM targets. Our method achieved higher average TMscore (0.740/0.754) than HHpred (0.665/0.709), CNFpred (0.674/0.705), CEthreader (0.687/0.721), and DeepThreader (0.698/0.725). The superiority of ProALIGN is more obvious for the TBM Hard and FM/TBM targets. In the case of TBM Hard targets, ProALIGN generated top1/top5 model with average TMscore of 0.688/0.703, which is significantly higher than HHpred (0.594/0.658), CNFpred (0.606/0.637), CEthreader (0.625/0.664) and DeepThreader (0.626/0.659). We can observe similar results when using GDT as quality measure.

**Table 5.** Threading performance on 80 CASP13 domain targets. Here, we show quality (measured using TMscore and GDT) of the top1/top5 predicted models

| Method | ALL |  | FM/TBM |  | TBM Hard |  | TBM Easy |  |
| --- | --- | --- | --- | --- | --- | --- | --- | --- |
|  | TMscore | GDT | TMscore | GDT | TMscore | GDT | TMscore | GDT |
| HHpred | 0.665/0.709 | 0.600/0.646 | 0.414/0.460 | 0.389/0.439 | 0.594/0.658 | 0.491/0.550 | 0.773/0.807 | 0.715/0.753 |
| CNFpred | 0.674/0.705 | 0.614/0.645 | 0.421/0.464 | 0.406/0.451 | 0.606/0.637 | 0.504/0.534 | 0.781/0.807 | 0.727/0.756 |
| CEthreader | 0.687/0.721 | 0.619/0.655 | 0.461/0.526 | 0.455/0.510 | 0.625/0.664 | 0.517/0.552 | 0.782/0.805 | 0.716/0.746 |
| DeepThreader | 0.698/0.725 | 0.636/0.663 | 0.503/0.532 | 0.475/0.512 | 0.626/0.659 | 0.528/0.555 | 0.790/0.813 | 0.735/0.760 |
| ProALIGN | **0.740/0.754** | **0.674/0.691** | **0.570/0.584** | **0.553/0.561** | **0.686/0.708** | **0.571/0.601** | **0.815/0.826** | **0.758/0.773** |

In addition to comparing the average TMscore of predicted models, we further examined the number of high-quality models. Head-to-head comparison of these approaches illustrated that ProALIGN generated more high-quality models than other approaches. For instance, ProALIGN outperformed CEthreader on 59 targets, while CEthreader outperformed ProALIGN on 20 targets. Moreover, on 17 targets, ProALIGN generated models with significantly higher quality (the difference of TMscore > 0.10). In addition, ProALIGN generated high-quality models (TMscore > 0.40) on 76 targets. These results clearly demonstrate the performance of alignment generation and alignment ranking used by ProALIGN.

### 3.6 Analysis of feature contributions to protein alignment

As mentioned above, ProALIGN uses a collection of protein features, including PSSM, secondary structure, solvent accessibility, and inter-residue distances. To access the contribution of these features to protein alignment, we fed various feature combination to the deep convolutional neural network used by ProALIGN.

Table 6 demonstrates that when only PSSM is used, ProALIGN showed an average TMscore of 0.518. The addition of secondary structure and solvent accessibility leads to increase of alignment accuracy (0.532). In contrast, the addition of inter-residue distance leads to more significant performance increase (0.545). These two types of features are complementary in certain degree, especially for the hard group; thus, when using all of these features, alignment accuracy further increases to 0.554.

**Table 6.** Reference-independent alignment accuracy of ProALIGN when using various combinations of features as input. Dataset: Test1K

| Group | #Alignments | PSSM |  | PSSM+SA+SS |  | PSSM+Dist |  | PSSM+SA+SS+Dist |  |
| --- | --- | --- | --- | --- | --- | --- | --- | --- | --- |
|  |  | TMscore | GDT | TMscore | GDT | TMscore | GDT | TMscore | GDT |
| Test1K Easy | 121 | 0.821 | 0.753 | 0.835 | 0.766 | 0.836 | 0.768 | 0.840 | 0.772 |
| Test1K Medium | 334 | 0.673 | 0.567 | 0.688 | 0.580 | 0.702 | 0.593 | 0.709 | 0.600 |
| Test1K Hard | 314 | 0.446 | 0.352 | 0.466 | 0.370 | 0.476 | 0.379 | 0.490 | 0.392 |
| Test1K | 769 | 0.604 | 0.508 | 0.620 | 0.524 | 0.631 | 0.533 | 0.640 | 0.542 |

### 3.7 The contribution of deep convolutional neural network to alignment likelihood inference

ProALIGN uses a deep convolutional neural network to infer alignment likelihood of all residue pairs in their entirety; thus, in principle, this network is expected to consider correlations among residue pairs and exploit characteristic pattern of alignment motifs. To examine this issue, we compared ProALIGN with another approach (denoted as IndivInferrer) that infers alignment likelihood for each residue pair individually using a fully-connected neural network.

Table 7 suggests that ProALIGN considerably outperformed IndivInferrer in terms of alignment accuracy (TMscore 0.640 vs. 0.602). We further performed a case study using 1qd1A as query protein and `3dfeA` as template. As shown in Figure 8, compared with ProALIGN, IndivInferrer reported more aligned regions with over-estimated likelihood. For example, IndivInferrer assigned the aligned regions (R80 - R135 of `1qd1A` vs. R60 - R100 of `3dfeA`, blue rectangle in Fig. 8(b)) with likelihood exceeding 0.5. These incorrect aligned regions precluded IndivInferrer from finding the optimal alignment. In fact, the alignment generated using IndivInferrer deviates greatly from the reference alignment. In contrast, ProALIGN over-estimated alignment likelihood on only a few regions. The alignment generated by ProALIGN is close to the reference alignment, and the model thus built is significantly better than that built using IndivInferrer (TMscore: 0.753 vs. 0.299).

**Fig. 8.**
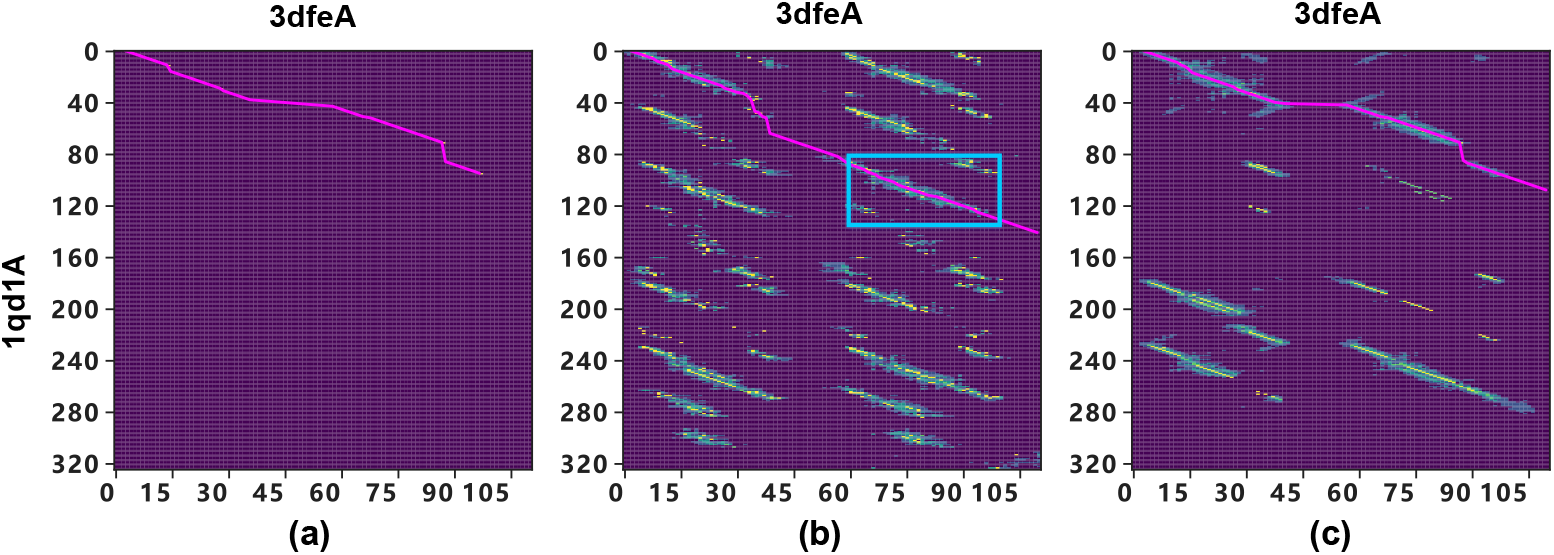
Comparison of ProALIGN and IndivInferrer in terms of accuracy of inferred alignment likelihood. (*a*) The reference alignment of proteins `1qd1A` and `3dfeA`. (*b*) Alignment likelihood inferred using IndivInferrer approach, which predict alignment likelihood for each residue pair individually. The aligned regions (E80-Q135 of `1qd1A` vs. G60-M100 of `3dfeA`, blue rectangle) is assigned with high likelihood exceeding 0.5. However, this aligned region is incorrect, and the derived alignment deviates greatly from the reference alignment. (*c*) Alignment likelihoods inferred using ProALIGN

**Table 7.** Reference-independent alignment accuracy of ProALIGN and IndivInferrer that infers alignment likelihood for each residue pair indvidually. Dataset: Test1K

| Group | #Alignments | IndivInferer |  | ProALIGN |  |
| --- | --- | --- | --- | --- | --- |
|  |  | TMscore | GDT | TMscore | GDT |
| Test1K Easy | 121 | 0.831 | 0.760 | 0.840 | 0.772 |
| Test1K Medium | 334 | 0.670 | 0.562 | 0.709 | 0.600 |
| Test1K Hard | 314 | 0.441 | 0.345 | 0.490 | 0.392 |
| Test1K | 769 | 0.602 | 0.505 | 0.640 | 0.542 |

We also observed that although ProALIGN reports multiple aligned regions with high likelihood, these regions essentially compose two alignments and both alignments can be used to build high-quality models (Fig. 9). In particular, using the first alignment (S1-R105 of `1qd1A` vs. S1-G111 of `3dfeA`), we obtained a predicted model with TMscore of 0.753, while using the second alignment (A172-C285 of `1qd1A` vs. S1-G111 of `3dfeA`), we could also obtain a perfect model with TMscore of 0.593.

**Fig. 9.**
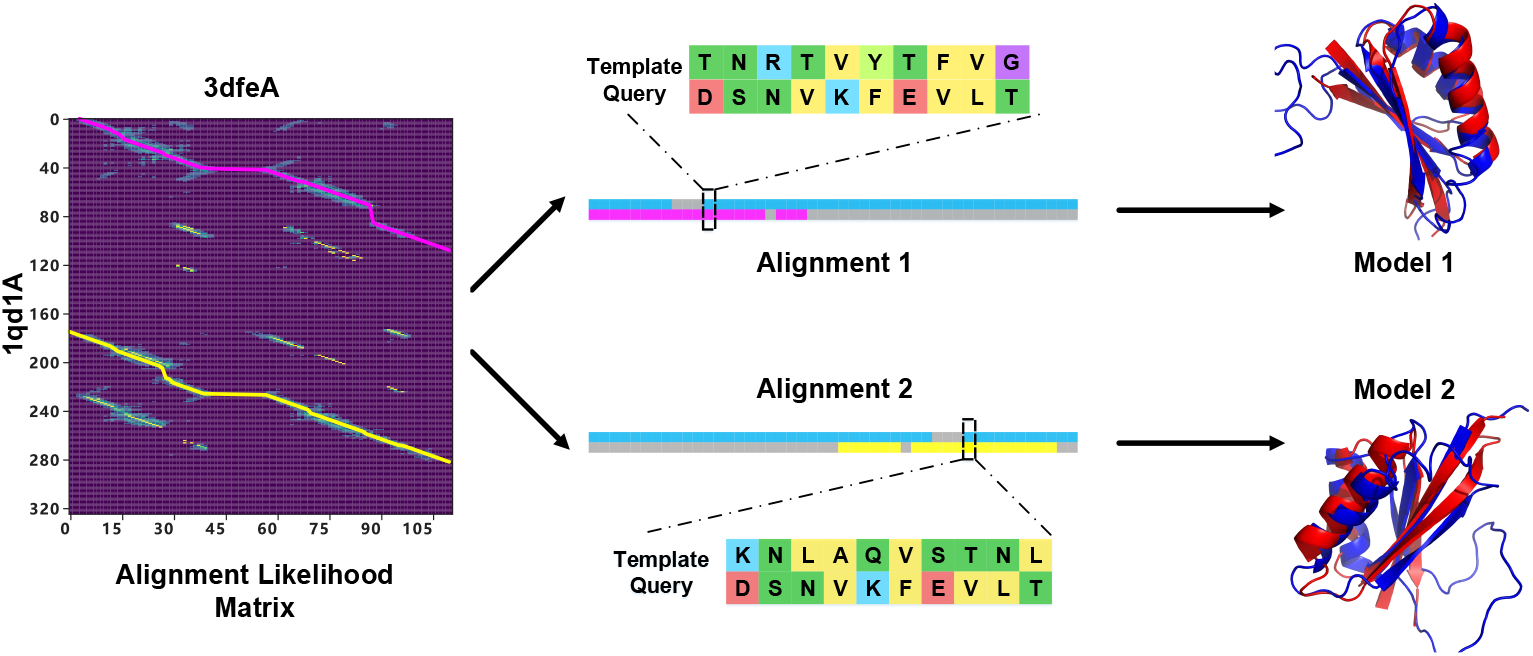
Two alignments constructed according to the alignment likelihoods inferred by ProALIGN for proteins `1qd1A` and `3dfeA`. Using the first alignment (magenta line, S1-R105 of `1qd1A` vs. S1-G111 of `3dfeA`), we obtained a predicted model with TMscore of 0.753, while using the second alignment (yellow line, A172-C285 of `1qd1A` vs. S1-G111 of `3dfeA`), we also obtained a perfect model with TMscore of 0.593.

Taken together, these results clearly demonstrate the advantage of deep convolutional network in considering correlations among residue pairs and the superiority of ProALIGN over IndivInferrer in inferring alignment likelihood.

## 4 Conclusion

The results presented in this study have clearly highlighted the special features of directly learning and inferring protein alignment through exploiting context-specific alignment motifs. The abilities of our approach to protein structure prediction have been demonstrated using 6,688 protein alignments and 80 CASP13 TBM targets as examples. Compared with the state-of-the-art threading approaches, ProALIGN could achieve much more accurate alignments and predicted structure models.

In principle, deep convolutional networks have large receptive fields when using deeper structure; however, the ability to detect correlation among long-range residues is limited in practice [17]. The application of graph convolutional networks[13] is promising in circumventing this limitation. In the future studies, this network will be incorporated into ProALIGN for the potential to model long-range correlations more accurately.

In summary, our approach ProALIGN should greatly facilitate understanding protein tertiary structures and functions.

## Acknowledgements

We greatly appreciate Shuai Cheng Li for fruitful discussions on this study.

