## Supplemental File 1 for "ProALIGN: Directly learning alignments for protein structure prediction via exploiting context-specific alignment motifs"

### 1 Relationship between ProALIGN's performance and quality of the input features

We examined the relationship between ProALIGN's performance and quality of the input features, including secondary structure, solvent accessibility, inter-residue distance, and MSA. As shown in Figure S1, there is no strong linear correlation between the quality of the predicted pair and the quality of the input features, which shows that ProALIGN does not rely solely on the quality of a certain feature. Relatively, the quality of MSA has the greatest influence with Pearson correlation coefficient 0.38.

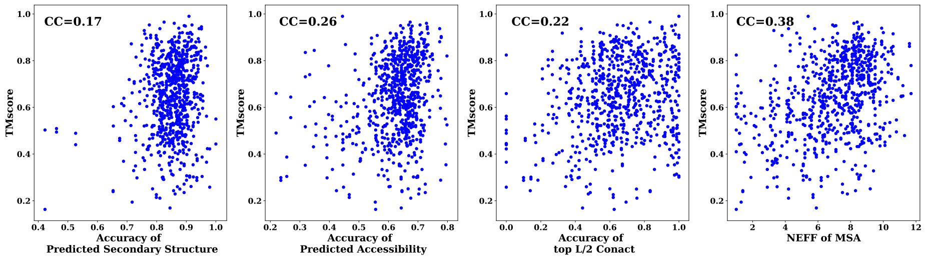


**Fig. S1.** Effects of the accuracy of input features on the final predicted model by ProALIGN. Dataset: Test1K. The input features include: (*a*) Predicted secondary structure, whose quality is measured based on the ratio of correct prediction by PSIPRED; (*b*) Solvent accessibility, whose quality is measured based on the ratio of correct prediction by RaptorX-property (*c*) Inter-residue distance, whose quality is measured using accuracy of the top L/2 predicted contacts; and (*d*) MSA, whose quality is measured using NEFF

### 2 The dynamic programming method for inferring alignment from alignment likelihood matrix

As described in Section 2.4, we calculated the optimal alignment $A_{opt}$ by maximizing likelihood of predicted matrix obtained by neural network, at the same time satisfying the definition of valid alignment, i.e.,

$$\arg\max_{A\in S} \log P(A|S,T) - \lambda \cdot r(A). \tag{1}$$

Here, $U$ represents the set of all valid alignments, and $r(A)$ represents a regular term to make the alignment compacted. In this study, we set $r(A)$ as the length of actual alignment area, i.e., the area from the first aligned residue pair to the last aligned residue pair in built alignment. Here we assume all elements in the predicted alignment likelihood matrix are independent of each other,

$$\arg\max_{A\in U} \log P(A|S,T) - \lambda \cdot r(A) \tag{2}$$

$$= \arg\max_{A\in U} \sum_{i}^{m} \sum_{j}^{n} \log P(A_{i,j}) - \lambda \cdot r(A) \tag{3}$$

, $r(A)m$ and $n$ represent the number of rows and columns of alignment likelihood matrix. We use dynamic programming to calculate the optimal alignment.

We use group of four symbols $< i,j,p,s >$ to describe the state in dynamic programming. Here, $i$ and $j$ represent the position of described state in alignment likelihood matrix. The symbol $p$ to describe whether matrix element $A_{ij}$ is in penalty status, in other word, whether in alignment penalty area. And then we further define three values of $p$: $L$, $I$ and $R$. $L$ represents $A_{ij}$ in the left area of alignment penalty area in alignment, $L$ represent $A_{ij}$ is in penalty area of alignment, and $R$ represent matrix element is in right area of alignment penalty area. In addition, we set another state $s$ to describe matrix element $A_{ij}$ choose $Y$ or $N$, where $Y$ represent match and $N$ represent not. We define dynamic programming matrices $M_{i,j,p,m}$ to represent the optimal score of the partial alignment which ends in alignment matrix element $A_{i,j}$ in penalty state $p$ and match state $s$. Here, the matrices are calculated recursively,

$$M_{i,j,p=L,s=N} = M_{i-1,j-1,p=L,s=N} + \sum_{k=1}^{i-1} \log P(A_{k,j-1} = 0) + \sum_{l=1}^{j-1} \log P(A_{i-1,l} = 0) + \log P(A_{i,j} = 0) \tag{4}$$

$$= \sum_{k=1}^{i} \sum_{l=1}^{j} \log P(A_{k,l} = 0) \tag{5}$$

$$M_{i,j,p=I,s=Y} = \begin{cases} M_{i-1,j-1,p=L,s=N} + \sum_{k=1}^{i-1} \log P(A_{k,j-1} = 0) + \sum_{l=1}^{j-1} \log P(A_{i-1,l} = 0) + \log P(A_{i,j} = 1) - \lambda \\ M_{i-1,j-1,p=I,s=Y} + \sum_{k=1}^{i-1} \log P(A_{k,j-1} = 0) + \sum_{l=1}^{j-1} \log P(A_{i-1,l} = 0) + \log P(A_{i,j} = 1) - \lambda \\ M_{i-1,j-1,p=I,s=N} + \sum_{k=1}^{i-1} \log P(A_{k,j-1} = 0) + \sum_{l=1}^{j-1} \log P(A_{i-1,l} = 0) + \log P(A_{i,j} = 1) - \lambda \end{cases} \tag{6}$$

$$M_{i,j,p=I,s=N} = \begin{cases} M_{i,j-1,p=I,s=N} + \sum_{k=1}^{i-1} \log P(A_{k,j-1} = 0) + \log P(A_{i,j} = 0) - \lambda \\ M_{i,j-1,p=I,s=Y} + \sum_{k=1}^{i-1} \log P(A_{k,j-1} = 0) + \log P(A_{i,j} = 0) - \lambda \\ M_{i-1,j,p=I,s=Y} + \sum_{l=1}^{j-1} \log P(A_{i-1,l} = 0) + \log P(A_{i,j} = 0) - \lambda \\ M_{i-1,j,p=I,s=N} + \sum_{l=1}^{j-1} \log P(A_{i-1,l} = 0) + \log P(A_{i,j} = 0) - \lambda \end{cases} \tag{7}$$

$$M_{i,j,p=R,s=N} = \begin{cases} M_{i,j-1,p=I,s=Y} + \sum_{k=1}^{i-1} \log P(A_{k,j-1} = 0) + \log P(A_{i,j} = 0) \\ M_{i-1,j,p=I,s=Y} + \sum_{l=1}^{j-1} \log P(A_{i-1,l} = 0) + \log P(A_{i,j} = 0) \\ M_{i,j-1,p=R,s=N} + \sum_{k=1}^{i-1} \log P(A_{k,j-1} = 0) + \log P(A_{i,j} = 0) \\ M_{i-1,j,p=R,s=N} + \sum_{l=1}^{j-1} \log P(A_{i-1,l} = 0) + \log P(A_{i,j} = 0) \end{cases} \tag{8}$$

### 3 Selection of hyper-parameter

Hyper-parameter $\lambda$ in dynamic programming method is determined by 1000 alignment pairs randomly selected from validation dataset. We tested 10 values of $\lambda$ for 0.1 to 1. The influence of different possible values are evaluated by the average predicted model quality (measured by TM-score). As shown in Table S1, ProALIGN is insensitive to the hyper-parameter $\lambda$, especially in the range from 0.1 to 0.4. Finally, we choose 0.3 as the final value.

**Table S1.** Influence of hyper-parameter. Dataset: Valid1K

| $\lambda$ | 0.1 | 0.2 | 0.3 | 0.4 | 0.5 | 0.6 | 0.7 | 0.8 | 0.9 | 1.0 |
|---|---|---|---|---|---|---|---|---|---|---|
| TMscore | 0.585 | 0.586 | 0.586 | 0.585 | 0.580 | 0.574 | 0.567 | 0.561 | 0.554 | 0.542 |

### 4 Alignment cases of failing in predicting high-quality alignment

There are still some cases when this method performed worse than HHPred or other existing methods. Here, we further analyze how these happened. In the case of Test1K dataset, we identified 6 template-query pairs, for which ProALIGN generated worse structural models than HHPred (TM-score difference $> 0.1$). These template-query pairs, together with failure reasons, are summarized as below.

As shown in the Table S2, the failures of ProALIGN can be divided into two cases: (1) Low-quality alignment likelihood matrix. In the cases of protein pair 3c12A-4g5aA, ProALIGN outputs

**Table S2.** Alignment Cases Worse than HHpred with TMscore > 0.1 . Dataset: Test1K

| Alignment | | NEFF | | Fold Type | | TMscore | | | |
|---|---|---|---|---|---|---|---|---|---|
| Template | Query | Template | Query | Template | Query | ProALIGN | HHpred | DeepAlign | Failure reason |
| 1yj7A | 1ctfA | 4.2 | 6.3 | a+b | a+b | 0.211 | 0.394 | 0.425 | Low-quality alignment likelihood matrix |
| 3c12A | 4g5aA | 7.4 | 1.3 | b | b | 0.194 | 0.451 | 0.465 | Low-quality alignment likelihood matrix |
| 3o0rB | 3dwwA | 7.8 | 6.3 | a | a | 0.480 | 0.585 | 0.683 | Low-quality alignment likelihood matrix |
| 4it7A | 2o9uX | 8.2 | 1.0 | a+b | a+b | 0.503 | 0.654 | 0.682 | Low-quality alignment likelihood matrix |
| 3kmiA | 1xl3C | 6.0 | 3.5 | a | a | 0.302 | 0.482 | 0.566 | Fail to construct alignment |
| 3hhmB | 1t3jA | 8.3 | 2.8 | a | a | 0.258 | 0.494 | 0.858 | Fail to construct alignment |

an alignment likelihood matrix (shown in panel C) that differs from the ideal structural alignment matrix reported by DeepAlign (shown in panel A), thus leading to the failure of ProALIGN.

(2) Fail to construct final alignment: In the case of protein pair 3hhmB-1t3jA, ProALIGN generated high-quality alignment likelihood matrix that is close to the ideal structural alignment matrix reported by DeepAlign. However, ProALIGN failed to construct the final alignment. The possible reason might be the insufficient penalty score for gaps in middle of alignment.

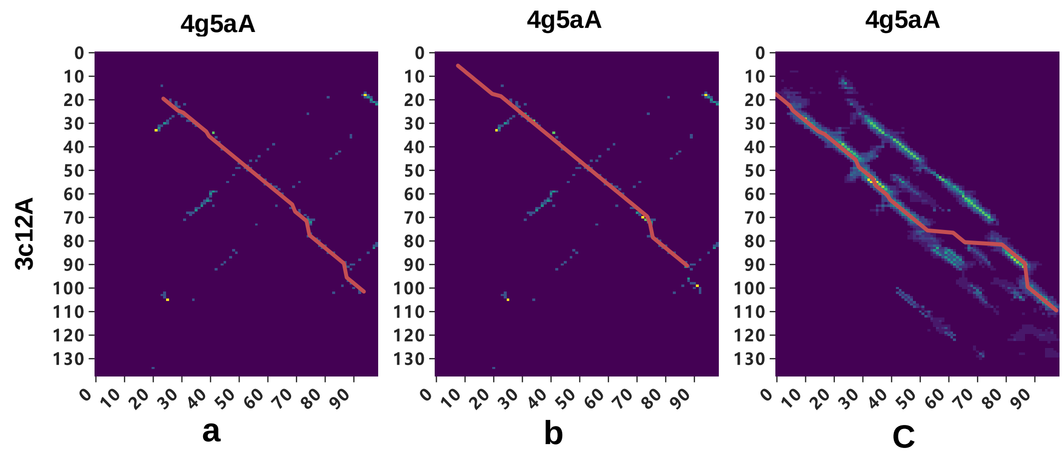


**Fig. S2.** Predicted alignment by HHPred and ProALIGN for proteins 3c12A-4g5aA. (a) Structural alignment matrix of the two proteins constructed by running DeepAlign. (b) Alignment predicted by HHpred (c) Predicted alignment likelihood matrix and the final predicted alignment

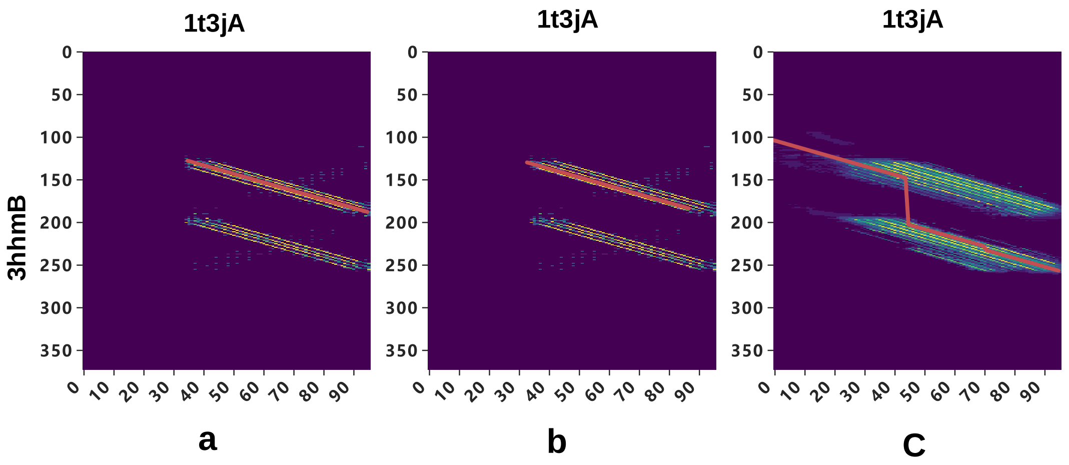


**Fig. S3.** Predicted alignment by HHPred and ProALIGN for proteins 3hhmB-1t3jA. (a) Structural alignment matrix of the two proteins constructed by running DeepAlign. (b) Alignment predicted by HHpred (c) Predicted alignment likelihood matrix and the final predicted alignment
